# Insular cortex projections to nucleus accumbens core mediate social approach to stressed juvenile rats

**DOI:** 10.1101/544221

**Authors:** Morgan M. Rogers-Carter, Anthony Djerdjaj, Katherine B. Gribbons, Juan A. Varela, John P. Christianson

**Author notes:** Corresponding Author: Morgan M. Rogers-Carter,. This manuscript was revised after peer review and includes new experiments and analysis. The manuscript was formatted to improve readability.

## Abstract

Social interactions are shaped by features of the interactants including age, emotion, sex and familiarity. Age-specific responses to social affect are evident when an adult male rat is presented with a pair of unfamiliar male conspecifics, one of which is stressed via 2 footshocks and the other naive to treatment. Adult test rats prefer to interact with stressed juvenile (PN30) conspecifics, but avoid stressed adult (PN50) conspecifics. This pattern depends upon the insular cortex (IC) which is anatomically connected to the nucleus accumbens core (NAc). The goal of this work was to test the necess ity of IC projections to NAc during social affective behavior. Here, bilateral pharmacological inhibition of the NAc with tetrodotoxin (1uM; 0.5ul/side) abolished the preference for stressed PN30, but did not alter interactions with PN50 conspecifics. Using a combination of retrograding tracing and c-Fos immunohistochemistry, we report that social interactions with stressed PN30 conspecifics elicit greater Fos immunoreactivity in IC → NAc neurons than interactions with naive PN30 conspecifics. Chemogenetic stimulation of IC terminals in the NAc increased social exploration with juvenile, but not adult, conspecifics, while chemogenetic inhibition of this tract blocked the preference to investigate stressed PN30 conspecifics, which expands upon our previous finding that optogenetic inhibition of IC projection neurons mediated approach and avoidance. These new findings suggest that outputs of IC to the NAc modulate social approach, which provides new insight to the neural circuitry underlying social decision-making.

**Significance Statement:** Social decision-making underlies an animal’s behavioral response to others in a range of social contexts. Previous findings indicate the insular cortex (IC) and the nucleus accumbens (NAc) play important roles in a range of social behaviors, and human neuroimaging implicates both IC and NAc in autism and other psychiatric disorders characterized by aberrant social cognition. To test whether IC projections to the NAc are involved in social decision making, circuit-specific chemogenetic manipulations demonstrated that the IC → NAc pathway mediates social approach toward distressed juvenile, but not adult, conspecifics. This finding is the first to implicate this circuit in rodent socioemotional behaviors and may be a neuroanatomical substrate for integration of emotion with social reward.

## Introduction

Social decision-making enables animals to respond to their social environments with flexible and context-appropriate behaviors (Insel & Fernald, 2004; O’Connell & Hofmann, 2011). An important feature of social decision-making is that animals appraise the affective state of others to generate behavioral responses. Accordingly, a number of studies report that rodents demonstrate either prosocial or avoidance behaviors toward distressed others (reviewed by (Meyza et al., 2017). These responses are modulated by situational factors, including familiarity, age, social rank, sex and prior experience (Li et al., 2014; Langford et al., 2006; Burkett et al., 2016; Rogers-Carter et al., 2018b; Rogers-Carter et al., 2018a; Kiyokawa et al., 2014; Ishii et al., 2012; Guzmán et al., 2009; Meyza et al., 2017), which suggest that socioemotional information is integrated with context and situational factors to inform behavioral actions. However, how areas in the brain that integrate affective cues, external stimuli and physiology can inform behavioral responses remains elusive.

Mechanistic studies have identified several structures in the social decision-making network (SDMN; (O’Connell & Hofmann, 2012) that produce flexible behavioral responses during social interactions (Hong et al., 2014; Yao et al., 2017; Lee et al., 2014; Yang & Tsai, 2017; Felix-Ortiz et al., 2016). The insular cortex (IC) is a site of multisensory integration (Rodgers et al., 2008; Gogolla et al., 2014; Gogolla, 2017) and it is highly interconnected with the SDMN positioning it as a hub for socioemotional information to be incorporated in social decision making processes (Rogers-Carter & Christianson, 2019). In addition to detecting social cues, social decision-making utilizes reward valuation to determine behavioral responses, and socioemotional information is likely one factor that influences reward valuation and behavior during healthy and disordered social cognition (Kohls et al., 2012). The nucleus accumbens core (NAc) is implicated in reward (Wise, 2002) and IC provides a major cortical input to the NAc (Wright & Groenewegen, 1996). Thus, there is compelling evidence to investigate the IC → NAc projection as a possible pathway by which socially relevant sensory information reaches the reward system to inform social decisions.

We have reported a role for insular cortex (IC) in a social affective preference (SAP) test in which rats demonstrate age-specific responses to stressed others (Rogers-Carter et al., 2018b). Specifically, in the SAP test, experimental rats exhibit approach toward stressed juvenile rats, but avoid exploring stressed adult rats. Optogenetic silencing of IC pyramidal neurons abolished this pattern. While the role of NAc in control of behavior during interactions with stressed individuals is unknown, it is well established that the nucleus accumbens (NAc) is critical to a number of social behaviors. NAc mediates social recognition (Ploeger et al., 1991) and reward with juvenile (Trezza et al., 2011) and adult conspecifics (Dölen et al., 2013). Regarding emotional cues, NAc encodes ultrasonic vocalization calls that convey stress (Willuhn et al., 2014) and shows enhanced dopaminergic transmission in subjects that observe a conspecific receiving footshock (Wu et al., 1999), which suggests the NAc encodes the socioemotional cues present during interactions with stressed others.

In the SAP paradigm, we hypothesize that IC is integral to the binding of social and situational factors to inform social decisions which, via glutamatergic efferents to nodes in the social decision making-network, can shape the pattern of circuit activity to favor approach or avoidant behavioral strategies. No prior mechanistic studies have explored the role of IC projection tracts in social behavior. Here we employed pharmacological and tract-specific manipulations to test if the IC → NAc projections are necessary for SAP behavior. We report that this tract is necessary and sufficient for approach toward stressed juvenile conspecifics, but not avoidance of stressed adult conspecifics.

## Materials and Methods

### Overview of Approach

To determine the role of the IC → NAc tract in social affective behavior we performed a series of experiments. First, to test the role of the NAc we used reversible inactivation by infusing tetrodotoxin (TTX) directly to NAc before SAP tests with either post-natal day 30 (PN30) juvenile or PN50 adult conspecifics. Next, we used retrograde tracing and Fos immunohistochemistry to quantify activity in IC → NAc neurons in experimental rats who underwent social interactions with either stressed or naïve juveniles. The cell counting results suggested that activity in this tract increases during encounters with stressed juveniles in the posterior IC. To establish the necessity and sufficiency of this tract to the social affective preference behavior we chose to use a tract-specific chemogenetic approach (Jaramillo et al., 2018b). This required that we first demonstrate feasibility of chemogenetic manipulations in the SAP test. To this end we virally expressed the inhibitory hM4Di receptor (Roth, 2016) in the IC and achieved inhibition via systemic administration of clozapine-n-oxide (CNO), which blocked the social preference for stressed juveniles and replicated our prior findings with pharmacological and optogenetic inhibition of IC (Rogers-Carter et al., 2018b). Following methods and considerations described by others (Mahler & Aston-Jones, 2018; Stachniak et al., 2014; Smith et al., 2016) to gain tract-specific, chemogenetic control of the IC → NAc circuit, we introduced hM4Di or hM3Dq in the IC under the synapsin promoter to achieve receptor expression in axon terminals in the NAc. CNO was delivered via bilateral cannula to the NAc, so that excitation or inhibition was selective to terminals of neurons that originate in IC, prior to testing.

To determine if IC → NAc neurons are necessary for social approach to stressed juveniles we infused CNO prior to SAP tests in rats expressing hM4Di in IC → NAc neurons. This interfered with approach directed at stressed juveniles but not adults. Because social interaction with a stressed juvenile increased Fos in IC → NAc neurons, and chemogenetic inhibition reduced social interaction, we hypothesized that a gain-of-function in the IC → NAc tract would be sufficient to promote social interaction. To test this, CNO was infused in experimental rats with hM3Dq in the IC → NAc tract prior to 1-on-1 social interactions with either naive juvenile or naive adult conspecifics. To test whether the IC contributes during social reward, per se, we adapted the SAP test to assess social novelty preference (Smith et al., 2018) in a task in which adult experimental rats preferred to investigate a novel socially rewarding adult conspecific compared to a familiar (cagemate) conspecific. Next, we inhibited the IC using hM4Di and systemic CNO and found that social novelty preference remained intact. A number of control, verification and chemogenetic specificity experiments are also described to address important concerns that relate to the interpretation of chemogenetic experiments.

### Subjects

Male Sprague-Dawley rats were purchased from Charles River Laboratories (Wilmington, MA) at either 250g, or postnatal (PN) day 22 or 42. All rats were acclimated to the vivarium in the Boston College Animal Care facility for a minimum of 7 days before surgery or behavioral testing resulting in groups of experimental adult rats at PN60-80 days, juvenile conspecifics at PN30 days and adult, postpubescent conspecifics at PN50 days at the time of testing. Subjects were housed in groups of 2 or 3, maintained on a 12h light/dark cycle, and provided food and water ad libitum. Behavioral testing occurred within the first 4 hours of the light cycle. All procedures were approved by the Boston College Institution Animal Care and Use Committee and adhered to the Guide for the Care and Use of Laboratory Animals.

### Social Affective Preference Tests

As previously reported (Rogers-Carter et al., 2018b), the SAP test allows for the quantification of social exploration directed at two novel conspecifics, one of which is stressed via footshock and the other naïve to stress treatment. On day 1, experimental rats were acclimated to a plastic tub cage arena (50 × 40 × 20cm; L × W × H) containing beta chip bedding and a wire lid. On day 2, the experimental rats were presented two naïve conspecifics, either juvenile (PN28-32) or adult (PN50-54) pairs, which were held in clear acrylic chambers (18 × 21 × 10cm; L × W × H). Conspecific chambers were comprised of acrylic rods spaced 1 cm apart center-to-center (Fig. 1C) and placed at opposing ends of the cage for a 5 min trial. The chambers were designed to be large enough to permit the contained rat to move freely. On days 3 and 4, one conspecific was stressed via 2 brief footshocks immediately prior to each test (5s duration, 1mA, 60s inter-shock-interval; Precision Regulated Animal Shocker, Coulbourn Instruments, Whitehall, PA). SAP tests began when the chambers containing naïve and stressed conspecifics were placed in the test arena. A trained observer blind to treatment quantified the amount of social exploration initiated by the experimental rat which included: nose-to-nose contact, nose-to-body contact or reaching towards the conspecifics. Social exploration was quantified first during live testing and again from digital video recordings by a second observer who was blind to experimental conditions to establish inter-rater-reliability. The correlation between observers for the SAP trials included in the current experiments was r_35_ = 0.922, R^2^ = 0.849, p < 0.0001.

**Figure 1.**
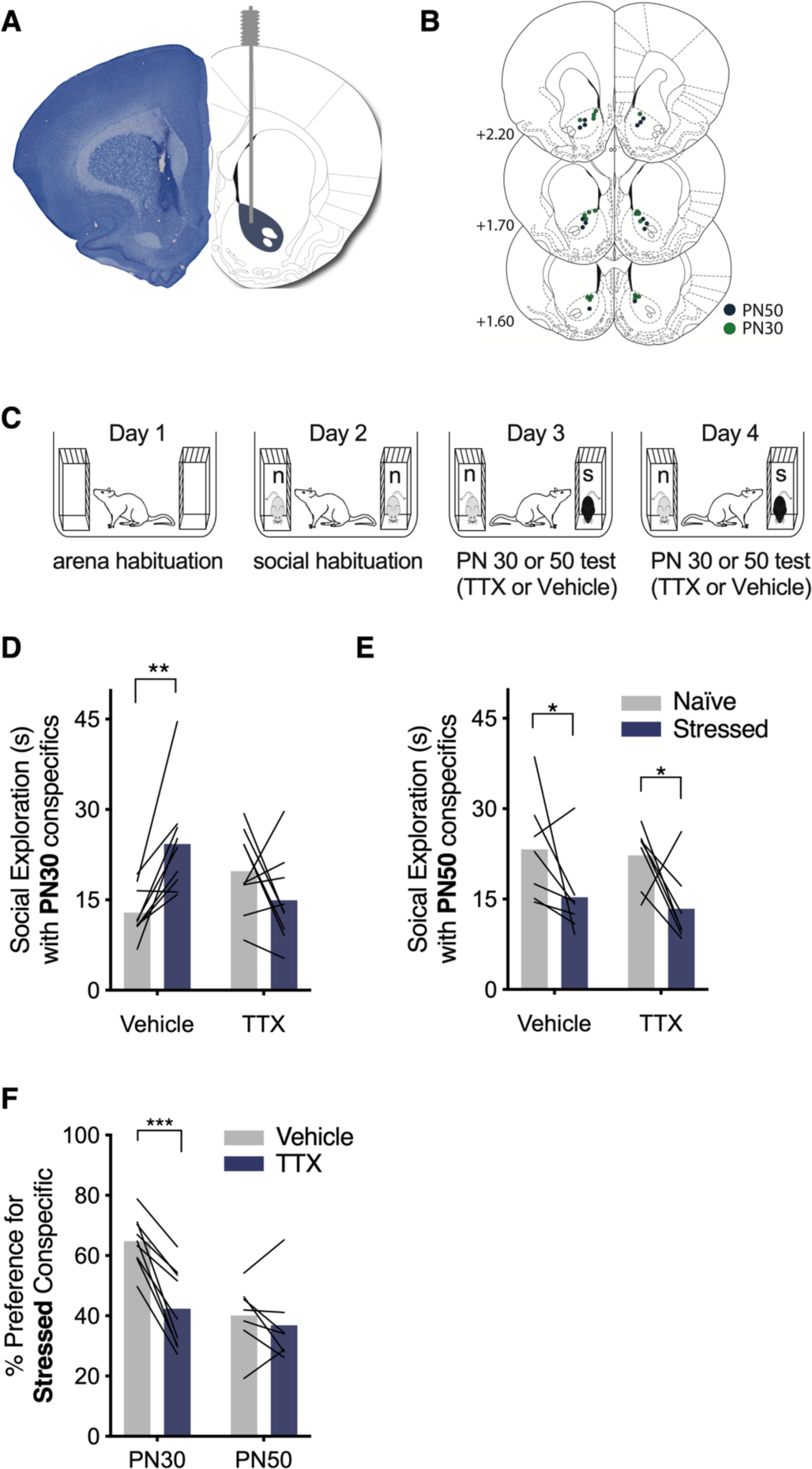
Pharmacological inhibition of the NAc abolished the social affective preference for stressed PN30, but not PN50, conspecifics. ***A***, Representative image of a cannula tract in NAc (left) and corresponding rat brain atlas diagram (right). ***B***, Map of bilateral cannula tip placements in NAc from experimental rats tested with PN30 (green; n = 9) and PN50 (blue; n = 7) conspecifics. ***C***, Diagram of SAP test procedure with naïve (n) and stressed (s) conspecifics. ***D***, Mean (with individual replicates) time spent exploring the naïve and stressed PN30 conspecifics during the 5 min trial. Vehicle-treated rats prefered to explore stressed PN30 conspecifics compared to naïve PN30 conspecifics (p = 0.006), which was abolished via bilateral infusion of tetrodotoxin (TTX; 100nM, 0.5µl/side) in NAc 15 min prior to testing (p = 0.203). ***E***, Mean (with individual replicates) time spent exploring the naïve and stressed PN50 conspecifics during the 5 min trial. Both vehicle (p = 0.031) and TTX-treated (p = 0.020) rats prefered to explore naïve PN50 conspecifics compared to the stressed PN50 conspecifics. ***F***, Mean (with individual replicates) data from (D,E) shown as the percentage of total social exploration time that was spent investigating the stressed conspecific. Rats tested with PN30 conspecifics show preference (as indicated by scores greater than 50%) for the stressed conspecific under vehicle treatment, which was significantly reduced after pharmacological inactivation of NAc with TTX (p = 0.0001). Rats tested with PN50 conspecifics show a preference for naïve conspecifics, which was unaffected by TTX treatment (p = 0.452). *p < 0.05, **p < 0.01, ***p < 0.001.

### Social Interaction

This procedure is used to quantify social exploration when an experimental test rat and a target conspecific were free to interact. On day 1, experimental rats were habituated to a standard plastic cage with beta chip bedding and wire lid for 1 h. On day 2, a juvenile or adult conspecific was introduced to the arena for 5 min and a trained observer quantified social interaction initiated by the experimental rat including pinning, sniffing and allogrooming. Exploration was quantified first during live testing and again by an observer blind to treatment from digital video recordings. The correlation between observers for the social interaction tests included in the current experiments was r_12_ = 0.997, R^2^=0.993, p < 0.0001.

### Cannula Placements and Virus Microinjections

Under inhaled isoflurane (2-5% v/v in O_2_), bilateral cannula (Plastics One) were implanted in the NAc (from Bregma: A/P +1.9mm, M/L +/- 1.8mm, D/V -6.5mm) and fixed in place with acrylic cement and stainless steel screws. For chemogenetic manipulations, 600nl of a viral vector containing either pAAV5-hSyn-hM4D(Gi)-mCherry (hM4Di; Addgene viral prep; Cat. No: 50475-AAV5; titer: 9×10^12^GC/mL), pAAV5- hSyn-hM3D(Gq)-mCherry (hM3Dq; Addgene viral prep; Cat. No: 50474-AAV5; titer: 4.8×10^12^GC/mL), or pAAV5-hSyn-mCherry (mCherry; Addgene viral prep; Cat. No: 114472), were microinjected bilaterally at 2 sites in the posterior insular cortex (from bregma: A/P –1.8mm and –1.6mm, M/L +/ – 6.5mm, D/V –6.9mm) at 100nl/minute and allowed 7 min for diffusion. These coordinates led to transduction within the posterior IC for consistency with our prior work and the results of the retrograde Fos counting experiment. In tract-specific chemogenetic manipulations, bilateral NAc cannula were implanted during the same surgery as described above. This approach allowed tract-specific control by directly infusing the hM3Dq and hM4Di agonist CNO to the terminals of IC neurons in NAc (see pharmacological manipulations). For retrograde tracing, 300nl of Cholera Toxin B conjugated to AlexaFluor 488 (CTb^488^, Thermo Fisher, Cat. No: C34775) were microinjected to the right hemisphere NAc as previously (Dong et al., 2017; Park et al., 2018) at a rate of 100nl/minute and allowed 4 min for diffusion. After surgery, rats were administered meloxicam (1mg/kg, Eloxiject, Henry Schein) and the antibiotic penicillin (12,000 Units, Combi-pen, Henry Schein) and allowed 2 to 3 weeks to recover before behavioral testing as described below.

### Pharmacological Manipulations

To reversibly inhibit NAc activity the voltage gated sodium channel blocker tetrodotoxin (TTX; Tocris Bioscience) was dissolved in sterile 0.9% saline to a concentration of 10µM. TTX or vehicle (sterile 0.9% saline) were infused bilaterally (0.5µl/side) to NAc 15 min prior to SAP testing (Christianson et al., 2011). For chemogenetic inhibition of IC cell bodies, experimental rats were weighed on the day of testing and given 0, 0.3 or 3mg/Kg CNO (Tocris Bioscience) dissolved in a vehicle of 10% DMSO and sterile 0.9% saline 45 min prior to SAP testing via i.p. injection. For tract-specific experiments, 1µM of CNO in a vehicle of 0.1% DMSO and sterile 0.9% saline was infused through the NAc guide cannula (0.5µl/side) 30 min prior to testing as in Mahler et al. (2014).

### Tissue Collection and Immunohistochemistry

Immediately after testing, rats from TTX experiments were overdosed with tribromoethanol (Fisher Scientific), decapitated, brains were dissected and flash frozen in 2-methylbutane (Fisher Scientific). All other rats were overdosed with tribromoethanol and perfused with cold 0.01M heparinized phosphate buffered saline (PBS) followed by 4% paraformaldehyde as in (Rogers-Carter et al., 2018b). Dissected brains were stored at 4°C in 4% paraformaldehyde for 24 h and transferred to 30% sucrose for 2 days. 40µm coronal sections were cut on a freezing cryostat (Leica). Alternating NAc sections were either directly mounted to gelatin subbed slides and stained with cresyl violet for cannula tip verification or stored in cryoprotectant as free-floating sections for immunostaining. IC sections were directly mounted to slides and coverslipped with Vectashield (Vector Labs) to confirm virus transduction. To visualize Fos or mCherry fibers in NAc, floating sections were quenched for endogenous enzymes with 3% H_2_0_2_ and blocked with 2% normal donkey serum (Jackson ImmunoResearch) in PBS-T (0.01% Triton-X 100) and then incubated overnight in rabbit anti-c-Fos primary antibody (1:5000; EMD millipore; Cat. No: ABE457; Lot: 2987437) or the rabbit anti-mCherry primary antibody (1:200; Invitrogen; Cat. No: PA5-34974; Lot: 114936) for 2 hr. After, sections were washed in PBS-T and incubated in biotinylated donkey-anti-rabbit secondary antibody (1:200; Jackson ImmunoResearch; Cat. No: 711-035-152) followed by the avidin-biotin complex kit (ABC Elite Kit, Vector Labs) and visualized with chromogen precipitate (NovaRed, Vector Labs). Stained sections were mounted on slides, dehydrated, cleared and coverslipped with Permount (Fisher Scientific). For Fos counting, tiled images of NAc tissue were acquired using a Zeiss AxioCam HRc digital camera through a 10x objective (N.A. 0.45) and Fos immunoreactive cells were manually counted in ImageJ by an observer blind to treatment. For tissue containing CTb^488^, floating sections were washed in PBS-T, blocked in 5% normal goat serum in PBS-T, and incubated overnight in rabbit-anti-c-Fos primary antibody as above at 4°C, washed, incubated in the Dylight 549 goat-anti-rabbit secondary antibody (1:500; Vector Labs; Cat. No: DI-1549), and floated on slides and coverslipped with Vectashield. For cell counting, green (CTb^488^) and red (Fos) channel tiled images of NAc were acquired using a Hamamatsu digital camera through a 10x objective (N.A. 0.45) and CTb^488^ and Fos-positive co-labeled cells were manually counted in ImageJ by a trained observer blind to treatment. Cells were counted within a 1250µm^2^ area in IC and from a 350 µm^2^ area in NAc.

### Electrophysiology

To verify the efficacy of hM4di and hM3Dq on neural activity whole cell recordings were made in 300µm thick coronal sections taken from a subset of rats at the conclusion of the behavioral experiments. Slice preparation, internal and external solutions, patch-clamp electrodes and current-clamp experiments and data analysis were conducted as previously (Rogers-Carter et al., 2018b). Briefly, whole cell recordings were achieved in cells visually identified as mCherry positive. Intrinsic excitability was established by quantifying the number of action potentials generated by a small depolarizing current injection (rheobase) in normal artificial cerebrospinal fluid, during application of CNO (10µM as in Venniro et al., 2017) and for an additional 10 min with CNO in the bath.

### Data analysis

The initial sample sizes for these experiments were determined by a priori power analysis (G*Power ver. 3.1) using the effect sizes from our prior observations using repeated measures designs. The effect size for data relating to the significant interaction in Figure 1C and Figure 2F of Rogers-Carter, Varela et al., 2018 were 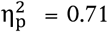 (N = 20, mixed model between and within subjects factors) and 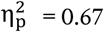 (N = 18, within subjects factors only), respectively. To achieve power = 0.80 to detect a similar large effect size for a 2-way interaction in a within-subjects model would require N = 8. Here we set out to meet or exceed these N by assigning a minimum of 8 subjects to each treatment condition. All data were included unless cannula implants or virus transduction missed the target site. Experimental treatments were counterbalanced in order and repeated measures designs were used except when noted. Social exploration times during SAP tests were compared using a 2-way ANOVA with conspecific stress (naïve or stressed) and drug treatment (vehicle or drug) analyzed as within-subjects factors. Data were visually inspected for normality and deemed suitable for parametric analysis; all data are depicted in the figures. To control for individual variation in sociality, exploration times were converted to a preference score, which was computed as the percentage of total social exploration time directed toward the stressed conspecific and analyzed with t-test or 2-way ANOVA, depending on experimental design. T-tests were used to compare social exploration in rats sacrificed for post mortem CTb, Fos, and CTb+Fos counts which were analyzed with mixed model ANOVAs with insula ROI as a repeated measure and conspecific stress as a between subject factor. All analyses were conducted in Prism 8.2 (GraphPad). Significant main effects and interactions were followed with Sidak post hoc comparisons to control experiment-wise type 1 errors to p < 0.05. Effect sizes are reported as η^2^ for t-tests or 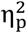 for main effects and interactions in ANOVAs.

**Figure 2.**
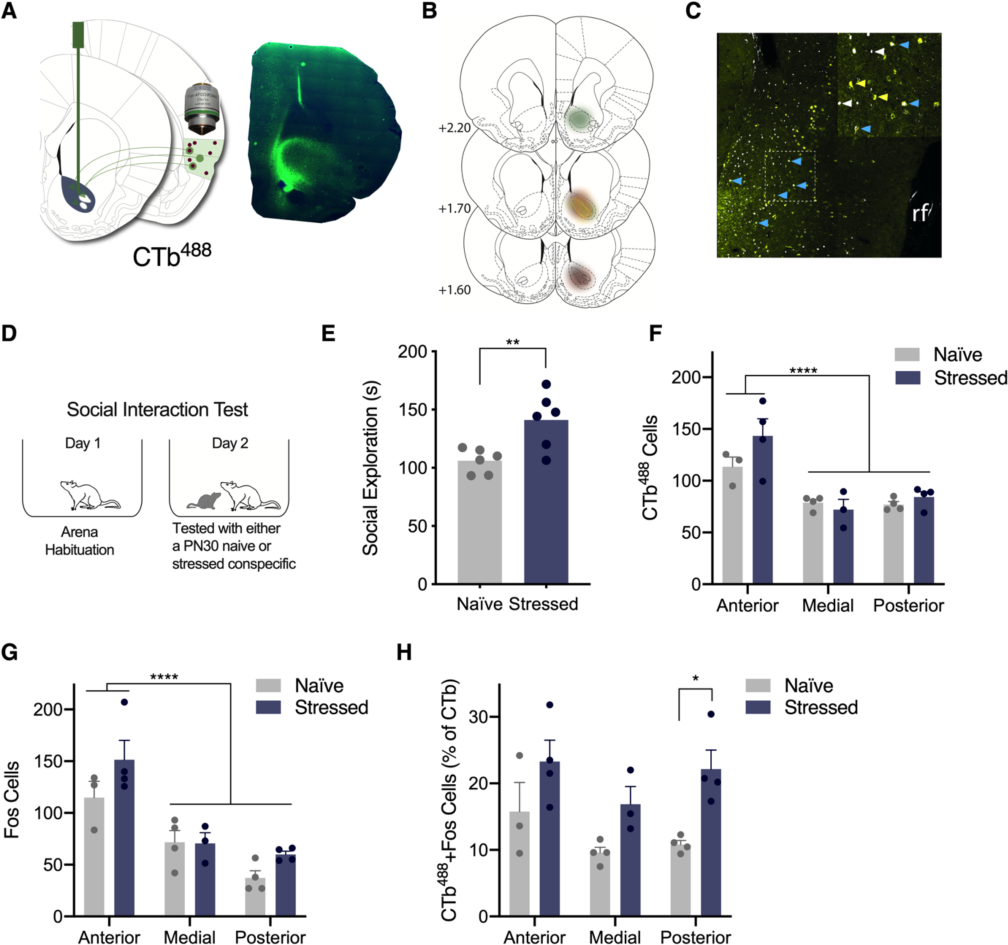
Exposure to stressed PN30 conspecifics elicited greater Fos immunoreactivity in IC → NAc neurons than exposure to naïve PN30 conspecifics. ***A***, Schematic depicting unilateral 300nl injections of the retrograde tracer CTb^488^ in the right hemisphere NAc (left) and representative image of CTb^488^ in NAc (right: green = CTb^488^, blue = DAPI). ***B***, CTb^488^ expression in NAc: each experimental rat is represented in a different color. ***C***, CTb^488^ and Fos-positive neurons in IC. Image is false colored; white = Fos, yellow = CTb^488^, blue = co-labeled cells. rf = rhinal fissure. ***D***, Diagram of behavioral procedures. 6 experimental rats explored naïve PN30 conspecifics and 6 experimental rats explored stressed PN30 conspecifics. ***E***, Mean (with individual values) social exploration time. Experimental rats presented stressed PN30 conspecifics engaged in more social exploration than rats presented naïve PN30 conspecifics (p = 0.008). ***F***, Counts of CTb^488^ positive cells in anterior, medial and posterior IC. More CTb^488^ positive cells were observed in anterior IC than medial or posterior IC (ps < 0.0001); there was no effect of conspecific stress on the number of CTb^488^-positive cells at any ROI. *G*, Counts of Fos positive cells in each IC region. More Fos positive cells were observed in anterior IC than medial or posterior (ps < 0.0001); there was no effect of conspecific stress on the number of Fos-positive cells at any ROI. *H*, Counts of colabeled CTb^488^ and Fos positive neurons. A main effect of Stress was observed (p = 0.0008), and experimental rats who explored stressed PN30 conspecifics expressed more Fos immunoreactivity in CTb^488^ positive neurons in posterior IC (p = 0.015) compared to experimental rats who explored naïve PN30 conspecifics. *p < 0.05, **p < 0.01, ****p < 0.0001

## Results

Pharmacological inactivation of NAc abolished preference for stressed PN30, but not PN50, conspecifics. To test if NAc was necessary for SAP behavior, experimental rats were implanted with bilateral NAc cannula (Fig. 1a and b), allowed 1 week of recovery, and underwent SAP testing with either PN30 (n = 9) or PN50 (n = 7) conspecifics. Experimental rats were tested after vehicle and drug injections on days 3 and 4 of testing and the amount of time the experimental rat explored each conspecific was recorded; treatment order was counterbalance (Fig. 1c). For rats tested with PN30 conspecifics, we report a significant Conspecific Stress X Drug interaction 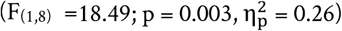. Following vehicle treatment, rats tested with PN30 conspecifics showed a preference to explore the stressed conspecific over the naïve (p = 0.006), which was abolished following bilateral pharmacological inactivation of NAc (p = 0.203, Fig. 1d). For rats tested with PN50 conspecifics, a 2-way ANOVA found a main effect of Conspecific Affect 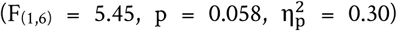 confirmed replication our previous findings (Rogers-Carter et al., 2018a; Rogers-Carter et al., 2018b) that experimental rats prefer to explore the naïve conspecific over the stressed following vehicle treatment (p = 0.031), which was also maintained after TTX treatment (p = 0.020; Fig. 1e). To compare the effect of TTX in NAc between age experiments, social exploration times were converted to percent preference scores and shown in Fig. 1f for comparison. Here, there were significant main effects of Conspecific Age 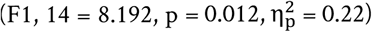, Drug (F1,14 = 25.31, p < 0.001, 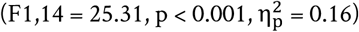) and a significant Conspecific Age by Drug interaction 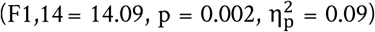 which reveals that preference scores from experimental rats tested with PN30, but not PN50 conspecifics, are sensitive to TTX treatment. Preference scores from experimental rats tested with PN30 conspecifics were significantly greater in vehicle trials than TTX trials (p < 0.0001). However, TTX did not alter preference scores in experimental rats tested with PN50 conspecifics (p = 0.651). This reflects that avoidance of stressed PN50 conspecifics was maintained after TTX treatment (Fig. 1f). These analyses demonstrate that the NAc is necessary for experimental rats’ preference to explore stressed PN30 but did not influence interactions with either stressed or naïve PN50, conspecifics.

Exposure to stressed PN30 conspecifics activated IC neurons that project to NAc. The NAc is a major target of IC afferents (Wright & Groenewegen, 1996) and the foregoing results suggest that these two structures are implicated in control of social responses to stressed juvenile conspecifics. Thus, we predicted that interaction with a stressed PN30 conspecific would activate NAc projecting IC neurons. To test this hypothesis, 12 experimental rats received unilateral microinjections of the retrograde tracer CTb^488^ in the left hemisphere NAc (Fig. 2a and b) as previously (Dong et al., 2017; Park et al., 2018) so that neurons that project from IC to NAc can be identified for cell counting (Fig. 2e). 2 weeks after surgery, in a between-subjects design, 6 experimental rats underwent social interactions with naïve PN30 conspecifics, and 6 underwent interactions with stressed PN30 conspecifics (Fig. 2c). As previously reported (Rogers-Carter et al., 2018b), experimental rats displayed higher levels of social exploration with stressed PN30 conspecifics compared to naïve PN30 conspecifics (t_(10)_ = 3.281, p = 0.008, η^2^ = 0.52; Fig. 2d). Rats were sacrificed 90 min after testing and tissue was processed and stained for fluorescent Fos expression. Tiled images of the IC ipsilateral to the NAc injection site were acquired at the following IC locations relative to Bregma: +2.52mm and 0mm (anterior IC), -0.24mm and -0.72mm (medial IC) and -1.72 and -2.04 (posterior IC) so that the number of Fos-positive, CTb^488^ -positive, and co-labeled neurons could be counted across the rostral-caudal extent of IC. All cells were counted in a field of veiw with a fixed area across all ROIs. Prior to conducting analysis, brains were screened for evidence of CTb^488^ deposits localized to the NAc (Fig. 2b). 4 animals, 2 from each stress group, did not exhibit CTb^488^ in the NAc and so were excluded from all subsequent analysis. We were unable to obtain counts from one subject from each treatment group at the anterior IC and medial IC regions resulting in ns = 3 or 4 observations per group. All individual replicates are depicted in Fig. 2f-h. To establish whether the resulting small sample size was sufficiently powered for our experimental questions, we used a mixed effects 2-way ANOVA with ROI and Conspecific Stress as within-subjects fators to compare CTb^488^ expression across anterior, medial and posterior IC. It is established that the insula projections to NAc is most dense in the anterior IC, compared to the posterior IC (Parks et al., 2015) and this trend was evident and supported by a significant main effect of ROI (F_(2,16)_ = 19.3, p < 0.0001). Anterior IC had significantly more CTb^488^ -positive cells than medial or posterior IC (ps < 0.001, Fig. 2f). Proceeding to the Fos analyses, a 2-way ANOVA revealed a main effect of ROI (F_(2,10)_ = 30.53, p < 0.0001). More Fos-positive cells were found in anterior IC, than medial or posterior IC (ps < 0.001, Fig. 2g) and there were no significant differences in Fos between conspecific stress groups. Regarding CTb^488^ and Fos-positive co-labeled neurons, we report a main effect of Conspecific Stress (F_(1,16)_ = 16.89, p = 0.0008). While the trend of greater CTb^488^ and Fos-positive was evident across ROIs, the only significant post-hoc comparision was observed between navie and stressed conspecific groups in the posterior IC (p = 0.015, Fig. 2h). In sum, exposure to stressed conspecifics led to greater activity in NAc-projecting IC neurons with the greatest differential activation observed in the posterior subregion of IC.

Chemogenetic inhibition of IC blocked social affective preference behavior. We previously reported (Rogers-Carter et al., 2018b) that the preference to explore stressed PN30 and naïve PN50 conspecifics was mediated by IC using an optogenetic approach. Here, an intersectional chemogenetic approach was preferred as the manipulation because it is less restrictive to movement, well suited to control neural activity in social behavior, and has been used to investigate IC projections to NAc and central amygdala in different paradigms (Jaramillo et al., 2018a; Venniro et al., 2017). Therefore, we first sought to replicate our prior finding and establish feasibility of chemogenetic control of IC neurons by using the inhibitory chemogenetic hM4Di receptor expressed in neurons under control of the synapsin promoter. Experimental rats (n = 10) were bilaterally transduced with pAAV5-hSyn-hM4D(Gi)-mCherry in IC (Fig. 3a and b). 2 weeks later, rats underwent the SAP test procedure with PN30 conspecifics following injections of vehicle or CNO (3mg/kg; i.p.), order counterbalanced, 45 min before testing (Fig. 3d and f). This produced a significant Conspecific Stress × Drug interaction 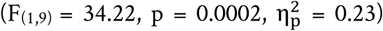. Vehicle-treated rats spent more time exploring stressed PN30 conspecifics than naïve PN30 conspecifics (p = 0.0001; Fig. 3f). This preference was abolished after inhibition of IC neurons as experimental rats tested with CNO did not show a preference for the naïve or stressed PN30 conspecific (p = 0.517; Fig. 3f). To interpret the effect of CNO in behavioral experiments a number of factors must be considered (MacLaren et al., 2016; Mahler & Aston-Jones, 2018; Manvich et al., 2018). To confirm these results were not an effect of CNO alone, experimental rats without virus transduction (n = 8, Fig. 3e) underwent the SAP procedure after i.p. CNO injections at the following doses: 0mg/kg, 0.3mg/kg, and 3.0mg/kg, the latter two of which have been previously reported to effectively modulate neurons transduced with chemogenetic viruses (Roth, 2016). A 2-way ANOVA with Conspecific Stress and Dose as within-subjects factors revealed a main effect of Conspecific Stress 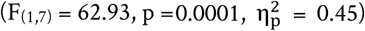. Post-hoc comparisons further revealed that, at each dose, experimental rats spent more time exploring the stressed PN30 conspecifics than the naïve (0mg/kg, p = 0.0009; 0.3mg/kg, p = 0.0036; 3.0mg/kg, p = 0.0063). These findings demonstrate that i.p. injections of CNO at 0.3 or 3.0mg/kg per se did not interfere with social affective preference behavior (Fig. 3g and h). To demonstrate virus efficacy, a cohort of experimental rats (n = 9) was bilaterally transduced with the sham virus pAAV5-hSyn-mCherry in IC (Fig. 3c), and two weeks later underwent SAP testing after vehicle and 3.0mg/kg i.p. CNO injections (order counterbalanced, Fig. 3e right side). Rats spent more time exploring the stressed PN30 conspecific after both vehicle (p = 0.003) and CNO (p = 0.002) treatment (Fig. 3f), which supports the main finding that IC inhibition achieved by chemogenetic inhibition blocked the social affective preference for the stressed PN30 conspecific and established feasibility for chemogenetic control of IC neurons during SAP testing. For side-by-side comparison of the effect of CNO in the sham and virus preparations, the data in Figs. 3e and f were converted to % preference scores and presented side by side for comparison in Fig. 3h. In no instance did administration of CNO alone (F_(7,14)_ = 0.343, p = 0.956, η^2^ < 0.01), or vehicle or CNO administration to rats with the sham mCherry virus (t_(8)_ = 0.010, p = 0.992, η^2^ < 0.01) disrupt the baseline preference to explore the stressed PN30 conspecific. In rats with transduced with hM4Di, administration of CNO attenuated the preference for the stressed conspecific (t_(9)_ = 5.654, p = 0.0003, η^2^= 0.78). Lastly, a subset of rats expressing hM4Di in the IC were sacrificed for acute slice whole cell recordings (n = 4 cells, Fig. 3h and i) as in (Rogers-Carter et al., 2018b). Following the methods described by Venniro et al., (2017) bath application of 10µM CNO reliably hyperpolarized and reduced excitability of IC neurons visually identified as mCherry positive at the time of recording. A one way repeated measures ANOVA revealed a main effect of CNO (F_(2,6)_ = 6.038, p = 0.040, η^2^= 0.66) and significantly fewer spikes under CNO compared to aCSF (p = 0.029).

**Figure 3.**
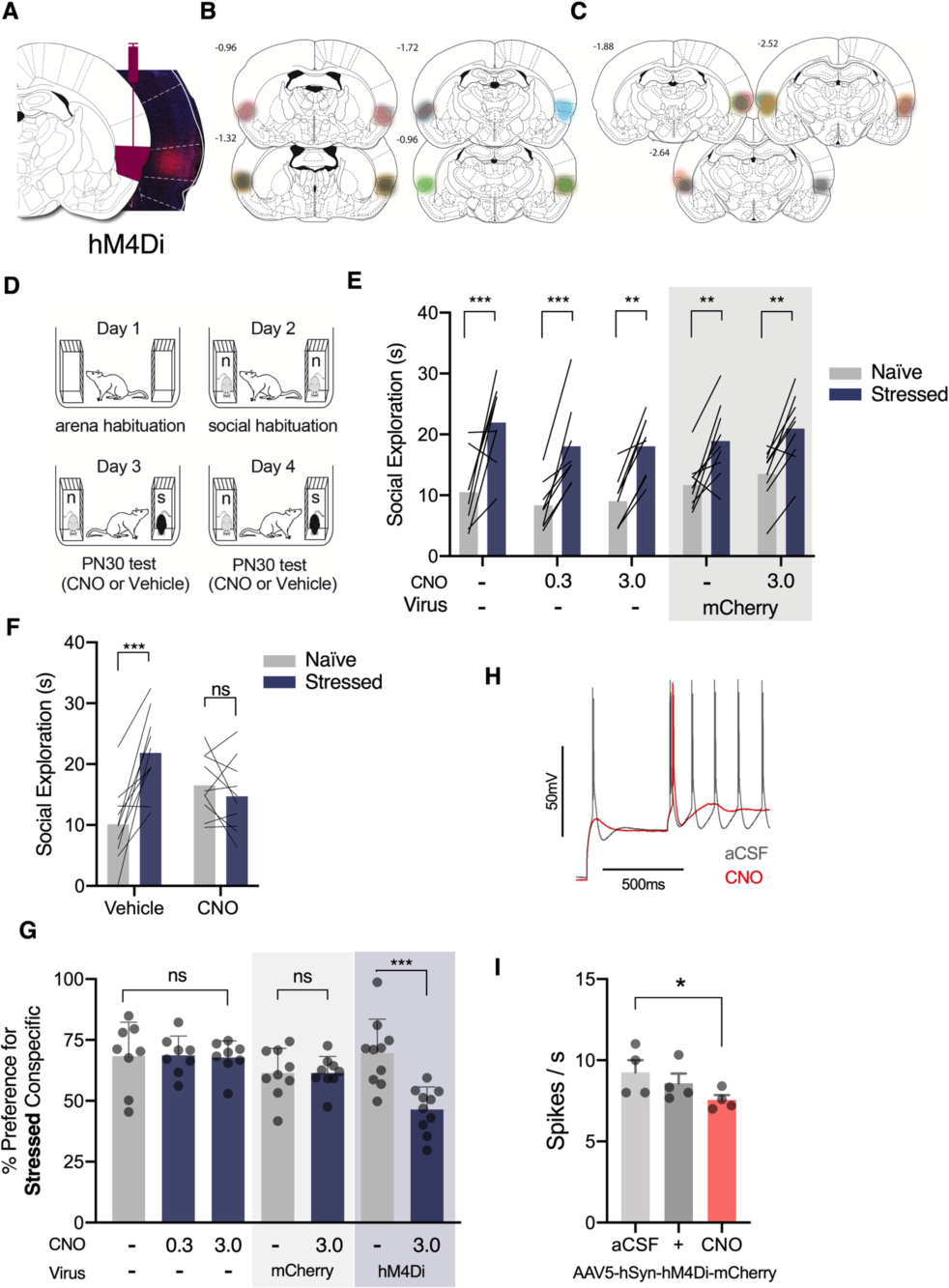
Chemogenetic inhibition of IC blocked the social affective preference for stressed PN30 conspecifics. ***A***, Representative image of pAAV5-hSyn-hM4D(Gi)-mCherry transduction in IC (right) and corresponding rat brain atlas diagram (left). ***B***, Map of bilateral IC pAAV5-hSyn-hM4D(Gi)-mCherry expression in experimental rats. Each subject is represented in a different color (n = 10); bold outlines represent the greatest intensity of mCherry expression and the corresponding faded overlay depicts the full extent of virus transduction. ***C***, Map of bilateral IC sham pAAV5-hSyn-mCherry expression, n = 8. ***D***, Diagram of SAP test procedure. ***E***, Control experiments: Experimental rats (n = 8) without viral transduction underwent the SAP test procedure 45 min after CNO injections at three doses: 0, 0.3 and 3.0mg/kg (i.p.). No dose of CNO disrupted the preference for the stressed PN30 conspecific (0mg/kg, p = 0.0009; 0.3mg/kg, p = 0.0036mg/kg; 3mg/kg, p = 0.0063). Experimental rats transduced with a sham virus (n = 9) preferred to explore the stressed PN30 conspecific under vehicle (p = 0.003) and 3mg/kg CNO (i.p.) (p = 0.002). ***F***, Mean (with individual replicates) time spent exploring the naïve and stressed PN30 conspecifics during the 5 min trial. Vehicle-treated rats spent more time exploring the stressed PN30 conspecifics than the naïve PN30 (p = 0.0001), which was abolished in trials following injections of CNO (3mg/kg; i.p.) 45 min before testing (p = 0.517). *G*, Data from (f) and (g) converted to % preference for the stressed conspecific. The only condition in which CNO reduced the preference for the stressed PN30 conspecific was in experimental rats transduced with hM4D(Gi) (p = 0.0003). *H*, Representative acute, whole-cell recording of hM4D(Gi) expressing IC neuron before and after bath application of CNO. *I*, After 10 min of bath application, CNO reduced the number of action potentials evoked by current injection in hM4D(Gi)-mCherry positive neurons in the IC. (+) indicates the beginning of CNO perfusion. *p < 0.5, **p < 0.01, ***p < 0.001.

Chemogenetic inhibition of IC terminals in NAc blocked the preference for the stressed PN30 conspecifics. To test if the IC → NAc pathway is necessary for SAP behavior with PN30 conspecifics, 10 experimental rats were transduced with the inhibitory hM4Di receptor via bilateral injection of pAAV5-hSyn-hM4D(Gi)-mCherry. Bilateral cannula were placed in NAc so that CNO could be directly administered to IC terminals (Fig. 4a left). Rats were only included in the final analysis if there was positive evidence of virus expression in IC as in Fig. 4a right; map of virus expression in Fig. 4b, bilateral cannula tip placement in NAc (Fig. 4a right, map in Fig. 4c), and mCherry immunoreactivity of putative IC axon terminals in NAc (Fig. 4a, right inset). Two experimental rats were excluded due to lack of virus expression. 3 weeks after testing, experimental rats underwent the SAP procedure with PN30 conspecifics after microinjections of vehicle or CNO in NAc 15 min before testing (order counterbalanced; Fig. 4d and e). A 2-way ANOVA of social exploration time revealed a main effect of Conspecific Stress 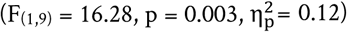; an interaction between Conspecific Stress and Drug approached significance 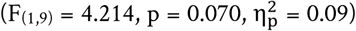.

**Figure 4.**
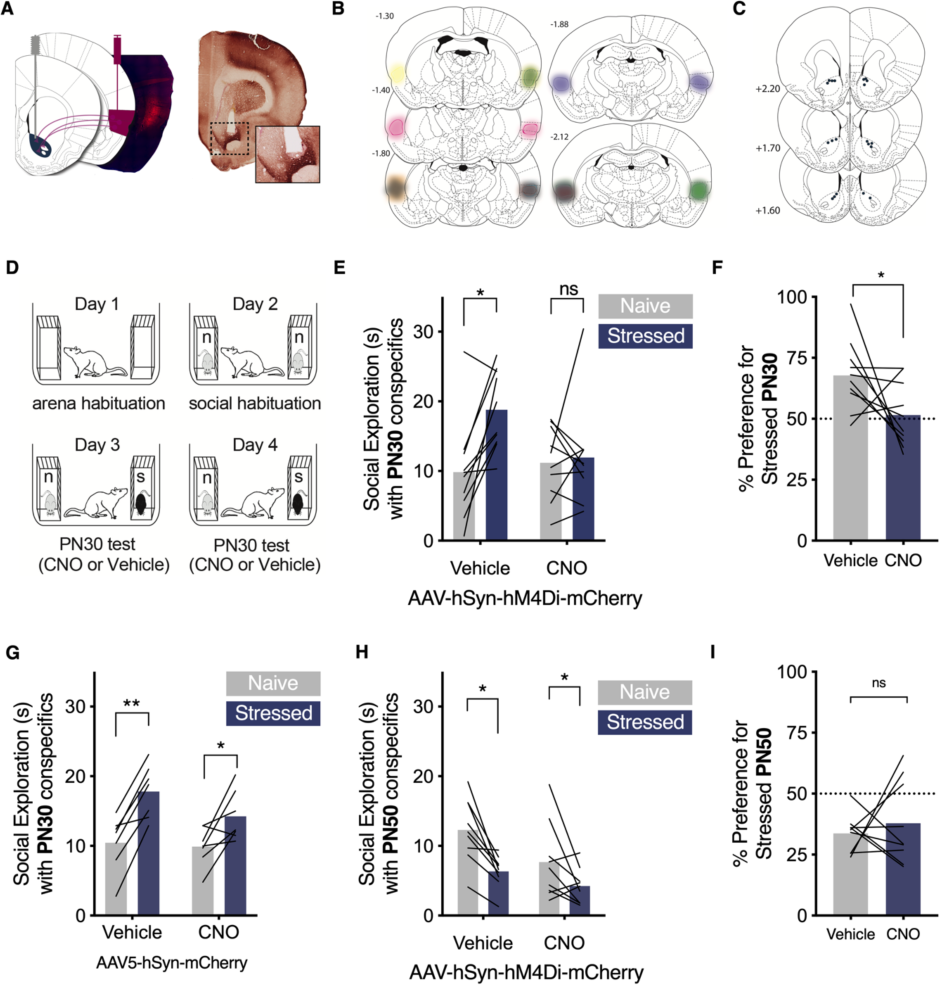
Chemogenetic inhibition of IC → NAc terminals blocked the social affective preference for stressed PN30 conspecifics. ***A***, Schematic of IC neurons transduced with pAAV5-hSyn-hM4D(Gi)-mCherry in IC and cannula implanted in NAc and representative image of virus transduction (left, red = native mCherry, blue = DAPI) and verification of cannula tip in NAc with terminal fiber expression (right, mCherry immunoreactivity). ***B***, Summary of mCherry expression in IC (n = 10); bold outlines represent the region of maximum mCherry expression and the corresponding faded overlay depicts the full extent of virus transduction. ***C***, Summary map of cannula tip placements in NAc. ***D***, Diagram of behavioral procedures; treatment order on Days 3 and 4 was counterbalanced. ***E***, Mean (with individual replicates, n = 10) time spent exploring the naïve and stressed PN30 conspecifics during the 5 min SAP test. Vehicle-treated rats spent more time exploring the stressed PN30 conspecifics than the naïve PN30 conspecifics (p = 0.023), which was abolished via 1µM CNO injections 45 min prior to testing through the guide cannula. ***F***, Data from (E) presented as the percent preference for the stressed conspecific. Experimental rats show a preference to explore the stressed conspecific under vehicle treatment, which was significantly reduced after pharmacological inhibition of IC terminals in NAc in CNO (p = 0.033). *G*, To control for non-specific effects of CNO in the NAc, a separate cohort of sham experimental rats without hM4D(Gi) transduction underwent the SAP test. Under vehicle treatment, rats preferred to explore the stressed PN30 conspecifics (p = 0.003), which was maintained after microinfusion of 1µM CNO to the NAc 45 min prior to testing (0.031). *H*, Mean (with individual replicates, n = 9) time spent exploring the naïve and stressed PN50 conspecifics during the 5 min SAP test after vehicle or 1µM CNO injections. Test rats spent more time exploring the naive conspecifics than the stressed PN50 conspecifics regardless of injection (main effect of Stress, p = 0.003). I, Data from (H) presented as the percent preference for the stressed conspecific; scores did not differ between vehicle and drug groups. * p < 0.05, **p < 0.01

In the vehicle condition, experimental rats preferred to explore the stressed PN30 conspecific over the naïve (p = 0.023). In the CNO condition, experimental rats did not exhibit preference for either the naïve or stressed conspecific (p = 0.959; Fig 4e). When social exploration was converted to percent preference scores (Fig. 4f), experimental rats showed scores that reflected more time spent with the stressed conspecific after vehicle injections, which was reduced by CNO 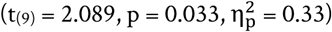. To rule out off-target effects of CNO, a separate cohort of experimental rats without any virus administration was bilaterally implanted with guide cannula in NAc and underwent SAP testing with PN30 conspecifics. One trial followed microinjections of vehicle solution, and the other followed microinjections of CNO. A 2- way ANOVA revealed a main effect of Drug 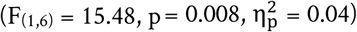 and Conspecific Stress 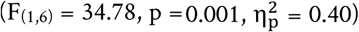. Experimental rats preferred to explore the stressed PN30 conspecific in both the vehicle (p = 0.003) and CNO (p = 0.031) conditions (Fig. 4g), which supports that the effect of CNO observed in hM4Di expressing rats cannot be attributed to off target effects of CNO on NAc function. To test whether IC → NAc neurons contribute to social avoidance of stressed adults, a separate cohort of rats received hM4D(Gi) injections in the IC and injection cannula placements in the NAc as above. Every aspect of the experiment was the same as in Figs. d and e except that the conspecifics were PN50 adult rats. Only rats meeting the criteria for IC hM4D(Gi) expression, bilateral NAc cannula placement, mCherry expression in NAc and displayed avoidance of the stressed conspecific under vehicle conditions were included in analysis. In adult SAP tests, test rats spent more time interacting with the naive conspecific after both vehicle and drug conditions. A 2 way ANOVA revealed a main effect of Conspecific Stress, 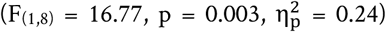. As in the CNO only expreiment, there was a small but significant effect of CNO on overal interaction levels evinced by a main effect of Drug 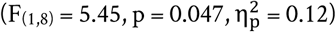. Converting these results to percent preference scores revealed no effect of CNO (t_(9)_= 0.57, p = 0.293). Consistent with the lack of effect of TTX in the NAc on adult SAP, chemogenetic inhibition of the IC → NAc terminals appears to be insufficient to alter preference for the niave rat in adult SAP tests. Together, these results indicate that IC projections to NAc are necessary to show a preference to explore the stressed PN30 conspecific compared to the naïve PN30.

Chemogenetic activation of IC terminals in NAc increased exploration with PN30, but not PN50, conspecifics. The results reviewed thus far suggest that during interactions with stressed juveniles IC projections to the NAc mediate the increase in social investigation directed toward the stressed targets. This set the stage to inquire whether this tract was sufficient to promote social exploration in the absence of social stress signals. We chose to again include PN50 conspecifics here because, while prior experiments suggested this pathway was not necessary for interactions with PN50 conspecifics, we did not have any prior data to rule out the possibility that the pathway was sufficient for interactions with PN50. To test this possibility, we augmented IC terminals in NAc by introducing the Gq coupled hM3Dq receptor via bilateral viral administration of pAAV5-hSyn-hM3D(Gq)-mCherry in IC in 10 experimental rats. Cannula were placed bilaterally in NAc so that CNO could be directly applied to IC terminals in NAc as above (Fig. 5a). For each experimental rat included in the analysis, we confirmed the virus transduction via fluorescent imaging of the mCherry reporter (Fig. 5a andc, right), the location of the cannula tips in NAc (Fig. 5b and c), and the presence of terminal fibers expressing mCherry in NAc (Fig. 5b, inset). Three rats were excluded due to misplaced cannula or lack of virus expression. 3 weeks after testing, experimental rats received social exploration tests over 5 days consisting of a habituation day, and then 2 tests with naïve PN30 conspecifics and 2 tests with naïve PN50 conspecifics. One trial for each conspecific age pairing occured after vehicle treatment, and the other following infusion of CNO (0.5µl/side) to NAc 30 min before testing (Fig. 5g). The order of conspecific age and order of drug treatment were counterbalanced which allowed for a within-subjects design. Social exploration was analyzed as a 2- way ANOVA with Conspecific Age and Drug as within-subjects factors 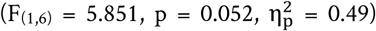. This analysis revealed main effects of Conspecific Age 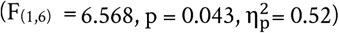 and Drug 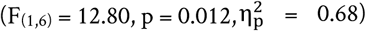. Post-hoc comparisons show mean social exploration was greater with PN30 conspecifics after CNO treatment than vehicle (p = 0.030), whereas mean social exploration with PN50 conspecifics did not differ between CNO and vehicle treatment (p = 0.998) (Fig. 5h). To verify hM3D(Gq) efficacy, a separate set of rats with hM3D(Gq) expression in the IC were used for acute whole-cell patch clamp recordings (Fig. 5f, n = 5 cells) which confirmed 10µM CNO reliably depolarized and inreased excitability of IC neurons visually identified as mCherry positive at the time of recording (Fig. 5g). A one-way repeated measures ANOVA revealed a main effect of CNO, (F_(2, 8)_ = 4.33, p = 0.053, η^2^ = 0.52) and significant post hoc comparison between aCSF and CNO, p = 0.032). Lastly, a day after testing ended, experimental rats received bilateral microinjections of vehicle (n = 4) or CNO (n = 3) 2h before sacrifice. NAc sections were stained for Fos to test if CNO was sufficient to increase activation of NAc neurons. CNO-treated rats expressed greater Fos-immunoreactivity compared to vehicle-treated rats (t_(5)_ = 2.727, p = 0.041, η^2^ = 0.60, Fig. 5h), supporting the efficacy of chemogenetic stimulation of IC-NAc terminals. These results indicate that excitation of IC terminals in NAc via CNO was sufficient to increase exploration with PN30, but not PN50 conspecifics.

**Figure 5.**
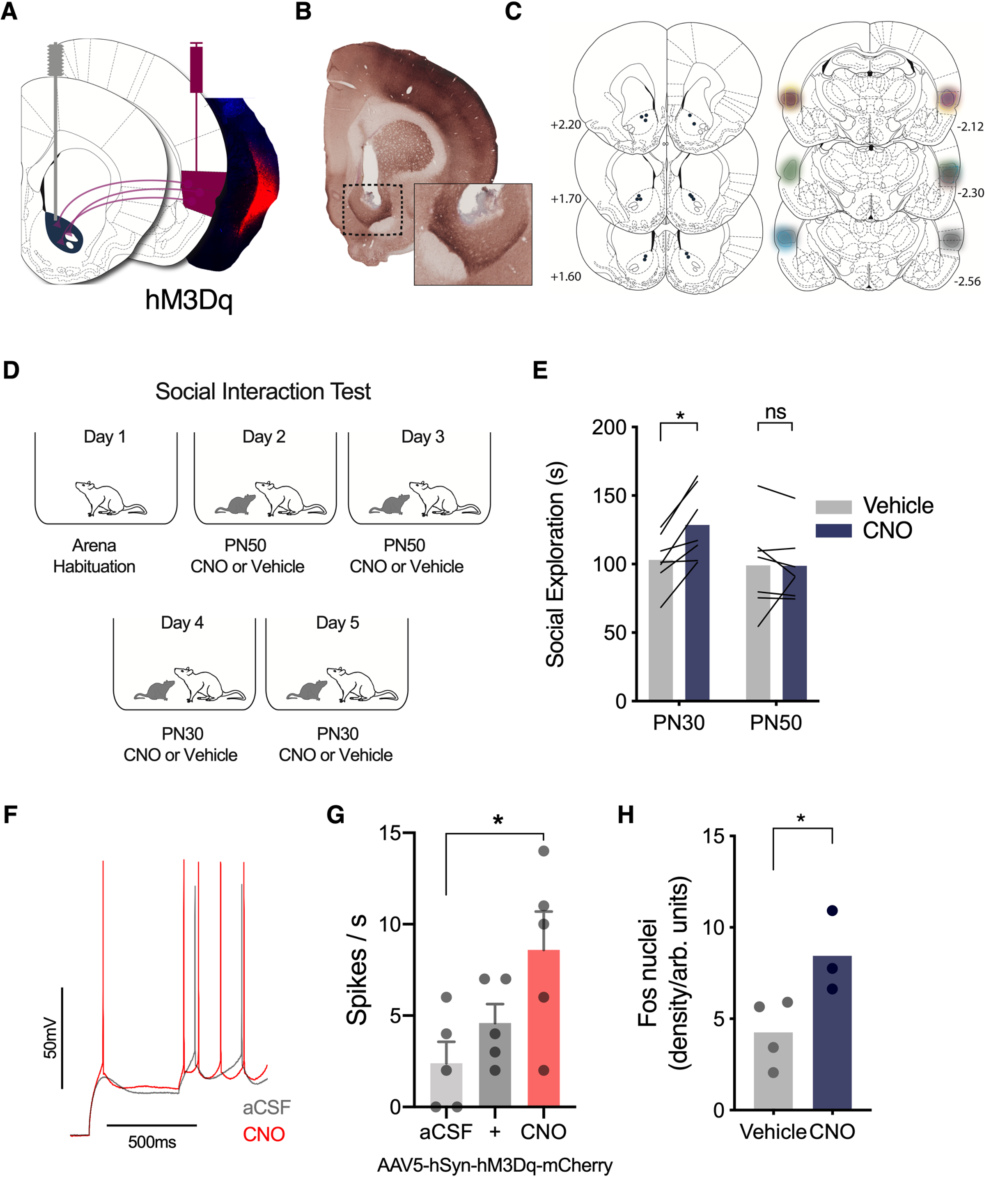
Chemogenetic excitation of IC terminals in NAc increased social exploration with PN30, but not PN50, conspecifics. ***A***, Schematic of IC neurons transduced with pAAV5-hSyn-hM3D(Gq)-mCherry in IC and cannula implanted in NAc (left). Terminal excitation of mCherry-positive fibers was achieved via bilateral clozapine-n-oxide (CNO) microinjections to NAc (1µM) 45 min before testing. Representative image of virus transduction (red = native mCherry expression, blue = DAPI) in IC (right). ***B***, Example of cannula tip placement in NAc and expression of terminal fibers in NAc (inset). ***C***, Map of cannula tip placements in NAc (left) and IC pAAV5-hSyn-hM3D(Gq)-mCherry expression in experimental rats (right). Each subject is represented by a different color (n = 7). ***D***, Diagram of social interaction trials. ***E***, Mean (with individual replicates) time spent exploring naïve PN30 and PN50 conspecifics following vehicle and CNO treatment (order counterbalanced). Experimental rats spent more time exploring naïve PN30 conspecifics following CNO injections than after vehicle treatment (p = 0.030), whereas social exploration with PN50 conspecifics was unaffected by drug treatment (p = 0.998). ***F***, Representative recording of CNO effect on hM3D(Gq) expressing neuron in whole cell recording. *G*, After 10 min of bath application, CNO increased the number of action potentials evoked by current injection in hM3Dq-mCherry positive neurons in the IC. (+) indicates the beginning of CNO perfusion. *H*, Greater NAc Fos immunoreactivity was observed in rats injected with CNO prior to perfusion, compared to rats treated with vehicle (p = 0.041). *p < 0.05.

Insular cortex is not necessary for social novelty preference. The foregoing results suggest the IC->NAc track is necessary and sufficient for social approach under conditions with a juvenile conspecific. Interestingly, manipulations of this tract did not influence behavior between two adult conspecifics. To establish whether IC is involved in situations of social reward per se we used the social novelty preference test in which others have shown that an experimental rat will spend more time exploring an unfamilair conspecific than a familiar animal and is dependent upon the NAc (Smith et al., 2018). We first adapted the social novelty preference test to be similar to the SAP test (Figure 6a). Experimental adult rats were housed in groups of 4 for 2 weeks. Social novelty testing occurred over 3 days; days 1 and 2 were habituation days exactly as in the SAP test. On Day 3, each experimental adult rat was presented with a pair of same age conspecifics, one of which was a cagemate (familiar condition) and the other was from another cage (unfamiliar) and time spent interacting with each was quantified (Figure 6b). Experimental rats (n = 10) spent significatnly more time exploring the unfamiliar conspecific 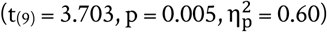. To test if preference for novelty is mediated by the IC, a separate set of experimental rats received hM4D(Gi) virus infusions to the IC exactly as in Fig. 2. After recovery, rats were habituated to the arena and social interaction and then received either CNO (3mg/Kg, i.p.) or vehicle injections 45 min before one of two social novelty preference tests (Days 3 and 4, drug order counterbalanced, Figure 6c). In this within-subjects design every combination of familiar and unfamiliar conspecifics was unique such that the experimental rat was never tested with the same individuals. Three rats were removed from analysis because of undetectable virus expression, or failure to show novelty preference under vehicle conditions; virus expression for the included rats is shown in Figure 6d. A repeated measures ANOVA revealed a main effect of Familiarity 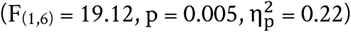; CNO had no effect on social novelty preference (p = 0.27).

**Figure 6.**
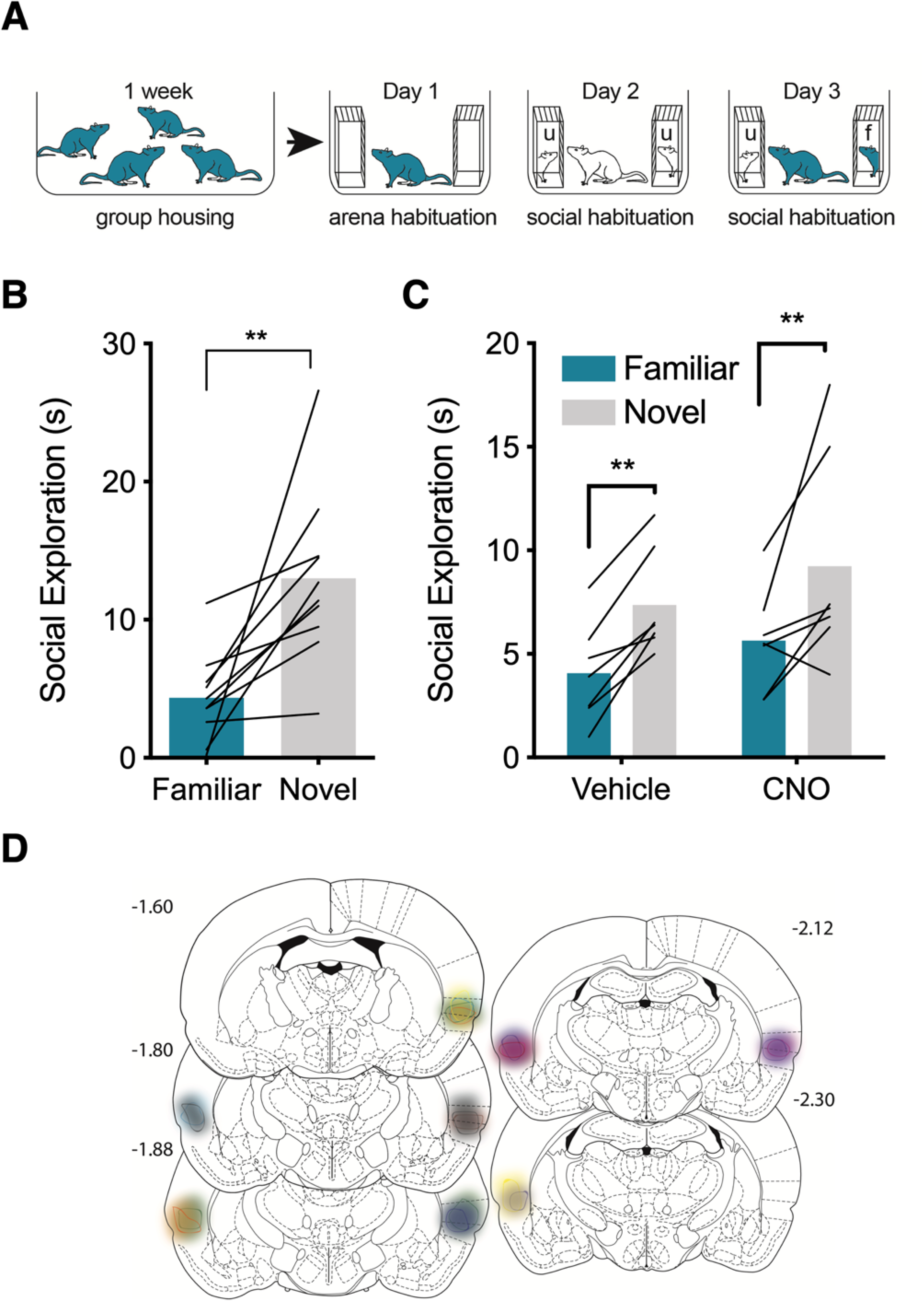
Insular cortex inhibition does not affect social novelty preference. ***A***, Experimetnal overview. Rats were housed in groups to establish familiarity. After 2 weeks, experimental rats were habituated to the test arena and social interaction then given a 5 minute novelty preference test with one of the experimental rat’s cagemates (f = familiar) and an unfamiliar rat (u = unfamiliar). ***B***, Mean (individual replicates) time spent in social interaction with the familiar and novel conspecifics. Experimental rats spent significantly more time investigating the unfamiliar target (**p = 0.005). ***C***, To test the role of IC in social novelty preference rats received pAAV5-hSyn-hM4D(Gi)-mCherry virus injections to IC and then housed in groups for at least 2 weeks. Social novelty preference tests were given 45 min after pretreatment with either vehicle or CNO (3mg/Kg, i.p.) on days 3 and 4. Experimental rats prefered interaction with the unfamiliar conspecific under both injection conditions (**main effect of Familiarity, p = 0.005). ***D***, Summary of mCherry expression indicating spread of hM4D(Gi) in the IC.

## Discussion

Here we investigated the role of the nucleus accumbens (NAc) and the projections from insular cortex (IC) to NAc in social affective behavior. In a social affective preference (SAP) test in which adult experimental rats demonstrate a preference to interact with stressed juvenile rats but avoid stressed adult rats (Rogers-Carter et al., 2018b), we observed effects of NAc and IC → NAc manipulations during interactions with juvenile conspecifics. First, pharmacological inactivation of the NAc prevented the expected preference for test rats to explore stressed PN30 conspecifics, but did not alter the avoidance of stressed PN50 conspecifics. Greater Fos immunoreactivity in NAc-projecting posterior IC neurons was evident in experimental rats following social interactions with stressed PN30 conspecifics compared to naïve PN30 conspecifics. In SAP tests, chemogenetic inhibition of this circuit blocked the preference for the stressed PN30 conspecific but had no apparent effect on avoidance of stressed PN50 conspecifics. Accordingly, chemogenetic stimulation of IC terminals in NAc increased exploration with PN30, but not PN50, conspecifics. Interestingly, chemogenetic inhibition of the IC did not alter social approach to a novel conspecific. These results implicate the IC → NAc pathway in prosocial behaviors directed toward juveniles. In light of our existing understanding of social decision-making, we suggest that the IC serves an important role in integrating sensory information about social stimuli with nodes in the social decision-making network, such as NAc, to modulate situationally appropriate social behaviors.

The involvement of NAc in social affective behavior is consistent with evidence implicating NAc involvement in myriad social processes. NAc mediates social recognition (Ploeger et al., 1991), social reward (Dölen et al., 2013; Smith et al., 2018), mating and sexual behavior (Xiao & Becker, 1997; Fujiwara & Chiba, 2018), social novelty-seeking (Smith et al., 2017), and interactions with both juvenile (Trezza et al., 2011) and adult (Dölen et al., 2013; Van den Berg et al., 1999) conspecifics. Furthermore, NAc is a node of the social decision-making network (O’Connell & Hofmann, 2011) and can decipher aversive from appetitive stimuli (Xiu et al., 2014) to modulate activity patterns in other social decision-making network structures based on the valence of sensory information (Johnson et al., 2017). In the SAP paradigm, the preference to explore stressed juveniles may be mediated by the NAc because this region integrates the valence of conspecific affect with reward valuation (Hamel et al., 2017). Consistent with this thought, when NAc was pharmacologically inactivated with TTX, rats did not show a social preference for the stressed PN30 conspecific and thus behaved as if each conspecific was equal in valence.

Like NAc, IC output neurons are necessary for the social affective preference for the stressed juvenile (Rogers-Carter et al., 2018b). IC is a site of multisensory integration (Rodgers et al., 2008; Gogolla et al., 2014; Gogolla, 2017) and is reciprocally connected to the extended amygdala (Shi & Cassell, 1998) which may underlie its known role in human emotional recognition (Adolphs et al., 2003; Gu et al., 2013; Terasawa et al., 2015). While the exact mechanism by which rodents convey emotion or affect is unknown, in the SAP test we reported that there are changes to several overt conspecific behaviors (e.g. self-grooming) and to the pattern of ultrasonic vocalizations which together could convey important features of conspecific stress and age (Rogers-Carter et al., 2018b). IC is anatomically situated to encode these features and relay this to the social decision making network. NAc is major target of IC efferents, and consistent with the hypothesis that IC projections to NAc mediate approach toward the stressed juvenile, we observed greater Fos IC → NAc neurons after interactions with stressed juveniles with the greatest difference observed in the posterior IC. A limitation of the current experiment is small sample size; this experiment may have been underpowered to detect smaller effects of conspecific stress on IC → NAc neurons in the anterior IC, which tend to be greater after interactions with stressed juveniles. Future studies are needed to directly address whether these projections are involved in social approach behavior. Further, using both inhibitory and excitatory tract-specific chemogenetic manipulations of IC → NAc terminals, we find that this tract is both necessary for the preference for the stressed juvenile, and sufficient to increase overall social exploration directed toward juvenile conspecifics. Together these findings demonstrate the modulatory role of IC → NAc projections on social approach in a preference paradigm.

Rats, like most mammals, require parental support to develop and survive (Lamers et al., 1986), and the same mechanisms that have evolved to enable animals to provide parental care may also underlie more complex social abilities (Scheiber et al., 2017; Marsh, 2018). Because these social actions are necessary for species survival, they are likely reinforced by the reward system in order to promote their occurrence (O’Connell & Hofmann, 2011). Although, we cannot be sure that approach to stressed juveniles is a prosocial reponse, increased NAc activity is associated with many prosocial behaviors. Optogenetic excitation of ventral tegmental neurons that project to the nucleus accumbens promoted mating behavior (Beny-Shefer et al., 2017) and microinfusion of a dopamine reuptake inhibitor increased social play in rats (Manduca et al., 2016). Similarly, oxytocin in NAc increases social approach (Steinman et al., 2018) and is important for forming pair bonds (Insel & Shapiro, 1992; Walum et al., 2012; Walum & Young, 2018). Our findings that chemogenetic manipulation of IC terminals in NAc relate to adult-to-juvenile behaviors adds to these NAc-dependent prosocial behaviors, and suggest an anatomical tract for socioemotional information to be integrated in social decision-making.

While the primary goal of this study was to investigate neural substrates of social behavior, this work adds to several studies which seek to identify functions of the IC using intersectional behavioral neuroscience methods. Recent reports in drug administration models show excitaiton of IC projections to central amygdala promote relapse of methamphetmaine use (Venniro et al., 2017) and IC projections to the bed nucleus of the stria terminalis mediate negative affect during alcohol abstinence (Centanni et al., 2019). Similarily, silencing of the IC projection to NAc decreased alcohol self-administration (Jaramillo et al., 2018b) and increased sensitivity to alcohol (Jaramillo et al., 2018a), and IC projections to the central amydala drive avoidance of an avertisve tastant (Schiff et al., 2018). While more work will add clarification, an emergent theme is that IC mediates both appetitive and aversive behavioral responses to external stimuli via different projections. IC contributes to both social approach and avoidance (Rogers-Carter et al., 2018b) the latter of which may be mediated IC projections to the central amygdala, which is necessary for social avoidance of stressed conspecifics (Ferretti et al., 2019).

An interesting issue concering the phenomena present in the SAP test is the motivation of experimental rats to approach stressed juveniles. Indeed, in much of the social behavior literature social stress signals lead to avaoidance. We speculated that adult-generated social affective cues were percieved as danger signals whereas juvenile emitted stress signals elicited stress buffering or parental behaviors from adults (Rogers-Carter et al., 2018a, Rogers-Carter et al., 2018b). The lack of IC involvement in the naturally rewarding social novelty preference test suggests that IC may have a specific role in adding multisensory or emotional content to the computations regarding social approach. The current findings parallel a number of important features of social behavior that are apparent in humans, namely that social stress cues modulate rodent social interactions in ways that mirror adult preference to lend help to juveniles while showing less empathy toward other adults (Batson & Powell, 2003). Abnormalities in social processes like emotion recognition are symptoms of several psychiatric conditions including autism spectrum disorders (Tracy et al., 2011; Lozier et al., 2014; Sato et al., 2017; Griffiths et al., 2017) and functional neuroimaging studies of individuals with autism find reduced insular activation to emotional stimuli (Di Martino et al., 2009; Hall et al., 2013; Morita et al., 2012), reduced striatal activation to social rewards (Schmitz et al., 2008; Scott-Van Zeeland et al., 2010; Supekar et al., 2018), and reduced functional connectivity between insula and ventral striatum (Fuccillo, 2016). The current experimental results provide important mechanistic support for this axis as a key to normative social cognition. Future research might take advantage of the SAP phenomenon to identify the tracts that mediate the avoidance of stressed adults and determine how genetic and experimental risk factors for autism spectrum disorders influence the IC and its targets in the social decision making network.

## Notes

Funding in support of this research was provided by the NSF Graduate Research Fellowship Program (M.M.R-C.); the Gianinno Family (J.P.C) and NIH Grant MH109545 (J.P.C.). We would like to thank Nancy McGilloway and the Boston College Animal Care Facility Staff for the husbandry of research animals involved in this study. pAAV-hSyn-hM3D(Gq)-mCherry and pAAV-hSyn-hM4D(Gi)-mCherry were gifts from Bryan Roth (Addgene plasmid # 50474; # 50475). pAAV-hSyn-mCherry was a gift from Karl Deisseroth (Addgene plasmid # 114472).

